# dSeqSb: A systems biology approach to decipher dynamics of host-pathogen interactions using temporal dual RNA-seq data

**DOI:** 10.1101/2022.02.28.482417

**Authors:** Mojdeh Dinarvand, Forrest Kock, Daniel Al Mouiee, Kaylee Vuong, Abhishek Vijayan, Afia Fariha Tanzim, AKM Azad, Anahit Penesyan, Natalia Castaño-Rodríguez, Fatemeh Vafaee

**Author notes:** Correspondence to: **Fatemeh Vafaee, PhD**, Deputy Director | UNSW Data Science Hub (uDASH), Team Leader | Artificial Intelligence in Biomedicine Laboratory, Senior Lecturer | Computational Biology, School of Biotechnology and Biomolecular Sciences, Room 2106, Level 2, E26, UNSW SYDNEY NSW 2052 AUSTRALIA, E, **Mojdeh Dinarvand, PhD**, School of Biotechnology and Biomolecular Sciences, Level 2, E26, UNSW SYDNEY NSW 2052 AUSTRALIA, E.

## Abstract

Infection triggers a dynamic cascade of reciprocal events between host and pathogen wherein the host activates complex mechanisms to recognise and kill pathogens while the pathogen adjusts its virulence and fitness to avoid eradication by the host. The interaction between the pathogen and the host results in large-scale changes in gene expression in both organisms. Dual RNA-seq, the simultaneous detection of host and pathogen transcripts, has become a leading approach to unravel complex molecular interactions between the host and the pathogen and is particularly informative for intracellular organisms. The amount of *in vitro* and *in vivo* dual RNA-seq data is rapidly growing which demands computational pipelines to effectively analyse such data. In particular, holistic, systems-level, and temporal analyses of dual RNA-seq data are essential to enable further insights into the host-pathogen transcriptional dynamics and potential interactions. Here, we developed an integrative network-driven bioinformatics pipeline, *dRNASb*, a systems biology-based computational pipeline to analyse temporal transcriptional clusters, incorporate molecular interaction networks (e.g., protein-protein interactions), identify topologically and functionally key transcripts in host and pathogen, and associate host and pathogen temporal transcriptome to decipher potential between-species interactions. The pipeline is applicable to various dual RNA-seq data from different species and experimental conditions. As a case study, we applied dRNASb to analyse temporal dual RNA-seq data of *Salmonella-infected* human cells, which enabled us to uncover genes contributing to the infection process and their potential functions and to identify potential host-pathogen interactions between host and pathogen genes. Overall, dRNASb has the potential to identify key genes involved in bacterial growth or host defence mechanisms for future uses as therapeutic targets.

## Introduction

Infectious diseases continue to be a major threat worldwide with devastating impacts on lives and economies around the globe. A better understanding of microbial infection mechanisms is central to the development of new antimicrobial therapies (1). During bacterial infections, a complex interplay is engaged by both bacterium and the host to negotiate the respective survival and defence strategies (2). Unravelling pathogen and host regulatory interactions, virulence mechanisms and innate responses, has led to the understanding of the dynamics of infectious processes and the development of therapeutics (3, 4). In studying infections, analysing transcriptomes of both organisms involved is key to better understand pathogenesis and disease progression (5, 6). Dual RNA sequencing (dRNA-seq) is an emerging powerful tool to capture whole transcriptomes, including coding and non-coding RNAs, of both host and pathogen simultaneously, in order to dissect the host-pathogen interplay and reveal the impacts that both organisms exert over each other during infection (7–10). Dual RNA-seq experiments are often designed to capture transcriptomics profiles of both organisms at different time-points after infection to enable the study of the molecular dynamics underlying host response and bacterial fitness (11). To understand the strategies employed by a pathogen for controlling or progressing of infection, a growing number of dRNA-seq experiments have been conducted under *in vivo* and *in vitro* conditions (2, 3, 12–14) for pathogens such as *Yersinia pseudotuberculosis* (15), *Pseudomonas aeruginosa* (16, 17), and *Mycobacterium leprae*(18), as well as intracellular bacteria such as *Mycobacterium tuberculosis* (7, 19) and *Piscirickettsia salmonis* (20).

Dual RNA-seq experiments inherently generate a mixture of transcripts from both host and pathogen requiring computational approaches to sort this mixture into the components associated with each organism. Multiple analytical workflows have been developed to align this read mixture to the reference genomes of the host and pathogen in model organisms (7, 9, 21) as well as non-model systems by leveraging the genomic resources of closely-related species (22). The majority of dRNA-seq bioinformatics pipelines generated to date have been focused on upstream analyses (e.g., quality control, alignment, read filtration, and normalisation) (22) with downstream analyses often limited to differential expression and functional analyses (7, 9, 21). Hence, there is a need for computational resources that offer more holistic and temporal downstream analyses of host and pathogen transcriptome enabling further insights into the host-pathogen transcriptional dynamics and potential interactions. In particular, systems biology approaches which integrate information on molecular interactions with gene expression information (23, 24), can provide a more holistic understanding of molecular intricacies underlying the infection process. However, the application of systems biology is still limited in this context.

To address this resource need, we developed an integrative network-driven bioinformatics pipeline, *dRNASb* (*<u>dRNA</u>-seq <u>S</u>ystems <u>b</u>iology-based analysis*), to analyse temporal transcriptional clusters, incorporate molecular interaction networks (e.g., protein-protein interactions), identify topologically and functionally key transcripts in host and pathogen, and associate host and pathogen temporal transcriptome to decipher potential between-species interactions. The pipeline is applicable to various dRNA-seq data from different species and experimental conditions. As a case study, we applied dRNASb to analyse dRNA-seq data of *Salmonella*-infected human cells (25) at six time-points post-infection (pi), which enabled us to uncover genes contributing to *Salmonella* infection and to identify their potential functions.

## Materials and Methods

### Data pre-processing and differential gene expression analysis

The pipeline accepts a dRNA-seq tab-delimited input file where read counts from both the host and the bacterium are collapsed into a single matrix of genes (rows) and time-points (columns). Rows with more than 50% missing values were filtered out. Raw counts were normalized using the trimmed mean of M-values (TMM), which corrects for differences in RNA composition and sample outliers while providing better comparability across-samples. Normalised data were then log2 transformed. Before differential expression analysis, reads were separated to the pathogen and host genome using the respective reference genomes (should be provided as input). Then, the *limma* package in R was used to identify differentially expressed (DE) genes, which uses linear modelling to describe expression data (26). The output is a list of DE genes compared with the baseline (i.e.,0 h). Genes with a false discovery rate (FDR) cut-off ≤ 0.05 (i.e., 5% false positives) and log fold change (LFC) of at least 2-fold (upregulation/downregulation), were considered as statistically significant DE genes. These lists can then be used as an input for the analysis of temporal gene expression profiles.

### Temporal analyses of gene expression profiles

To unravel transcriptional dynamics, fuzzy clustering was conducted on DE genes in at least one time-point. We used Mfuzz package in R (27) which is developed for noise-robust soft clustering of gene expression time-series data. The number of clusters should be first identified for which we performed repeated soft clustering for a range of cluster numbers using ‘*Dmir*’ function and reported the minimum centroid distance (MCD), (i.e., the minimum distance between two cluster centres). The optimal cluster number was then determined by plotting MCD against cluster numbers and selecting a number where there is a drop of MCD and slower decrease afterwards (*a.k.a*. the elbow rule). After soft clustering, each gene is assigned a membership score representing the degree of its association to each cluster that reflects a particular temporal pattern. The objective of Mfuzz is to compute membership values to minimize the within cluster distances and maximize the between-cluster distances. The output of Mfuzz is a membership list for each gene class. The membership value ranges from 0 to 1, with higher values indicating that a gene is more likely to belong to a particular class.

### Pathway enrichment analyses for host and pathogen

To identify potential functions associated with DE genes in each temporal cluster, the Cluster Evaluation R (clueR) package in R (28) was used to assess the enrichment of different functional terms by DE genes in each cluster, using Fisher’s Exact test with hypergeometric null distribution to calculate the probability of pathways associated with a set of DE genes within a cluster; adjusted *P<0.05* was considered to indicate statistically significant associations. Enrichment analysis relies on a reference dataset that annotates genes based on associated pathways or other functional terms. A database of pathways, regulons, and genomic islands was constructed using information obtained from the BioCyc.org Database and relevant literature sources (29) for the bacterium (i.e., *Salmonella enterica SL1344*). The BioCyc.org is unique in integrating a diverse range of data and providing a high level of curation for important microbes as well as model eukaryotic organisms and Homo sapiens (30). The Kyoto Encyclopedia of Genes and Genomes (KEGG) and Gene Ontology Consortium (Amigo: http://amigo.geneontology.org) were used to retrieve functional annotations for the host (i.e., human).

### Gene Ontology enrichment analysis and visualization for host and pathogen

Gene Ontology (GO) is an annotation database structured as a hierarchical directed acyclic graph where each node represents a class of gene function (GO term), and the connection between two GO terms indicates different relationships, such as “is a” or “part of”. GO terms are categorised into biological processes (BPs), molecular functions (MFs) and cellular components (CCs) (31). Statistical enrichment of GO terms was conducted using Enrichr tool which comprise an up-to-date libraries of diverse gene sets including gene ontology BP, MF and CC gene sets.(32) Similarly, statistical overrepresentation analyses were performed using Fisher’s exact test with hypergeometric null distribution. Visualization of enrichment was accomplished via bar plots and bubble plots using R scripts.

### Protein-protein interaction network construction, analyses, and visualization

A protein-protein interaction (PPI) databases for the pathogen (e.g., SalmoNet, salmonet.org (33) for the pathogen of interest in this study) and host (e.g., human STRING, string-db.org (34)) were retrieved and the corresponding network was constructed using *igraph* package (35) for network topology analyses. Accordingly, multiple network measures including node degrees, betweenness centrality (36), closeness centrality (36) and modularity (via community detection) were identified. The functions of the hub genes (i.e., those genes with high degree of connectivity), were identified as described before. Community structure detection was constructed using cluster edge betweenness function in the *igraph* package to cluster similar groups of hub genes (i.e. those which have the highest similarity degree) (37).

Inter and intraspecies correlation analysis of pathogen and host transcriptome Pearson correlation coefficients were calculated between and across host and pathogen DE genes, and *P values* were calculated using the function *cor.test* in R. To account for a possible temporal delay between *Salmonella* expression changes and effect manifestation in the host cell, a time-shift was allowed. This means the expressions of pathogen genes at each time-point were compared to the host expression at the subsequent time point. Once the correlation network was built (correlation coefficient < −0.7 or > 0.7, and *adjusted P < 0.05*), densely connected genes were identified via Louvain clustering, which can quickly find clusters with high modularity in large network (38). Host and pathogen genes within each cluster were further investigated to identify genes potentially important during early stage of infection.

## Results and Discussion

The dRNASb bioinformatics pipeline comprises the following steps: filtering and normalisation of count data, identification of DE genes in the host and pathogen, temporal gene expression clustering, functional enrichment analyses, network analysis, visualization, and inter/intraspecies correlation analyses (Figure 1). The pipeline, as mentioned above, is coded in R (version-4.0). The model system used in this work to study the bioinformatic pipeline comprised human cervix carcinoma cells (HeLa-S3; ATCC CCL-2.2) (host) infected with *Salmonella enterica* serovar Typhimurium strain SL1344 (bacterium), which is a well-defined host–bacteria system (39). The pipeline however can be repurposed to analyse temporal dRNA-seq data from other host-bacteria systems of interest.

**Figure 1.**
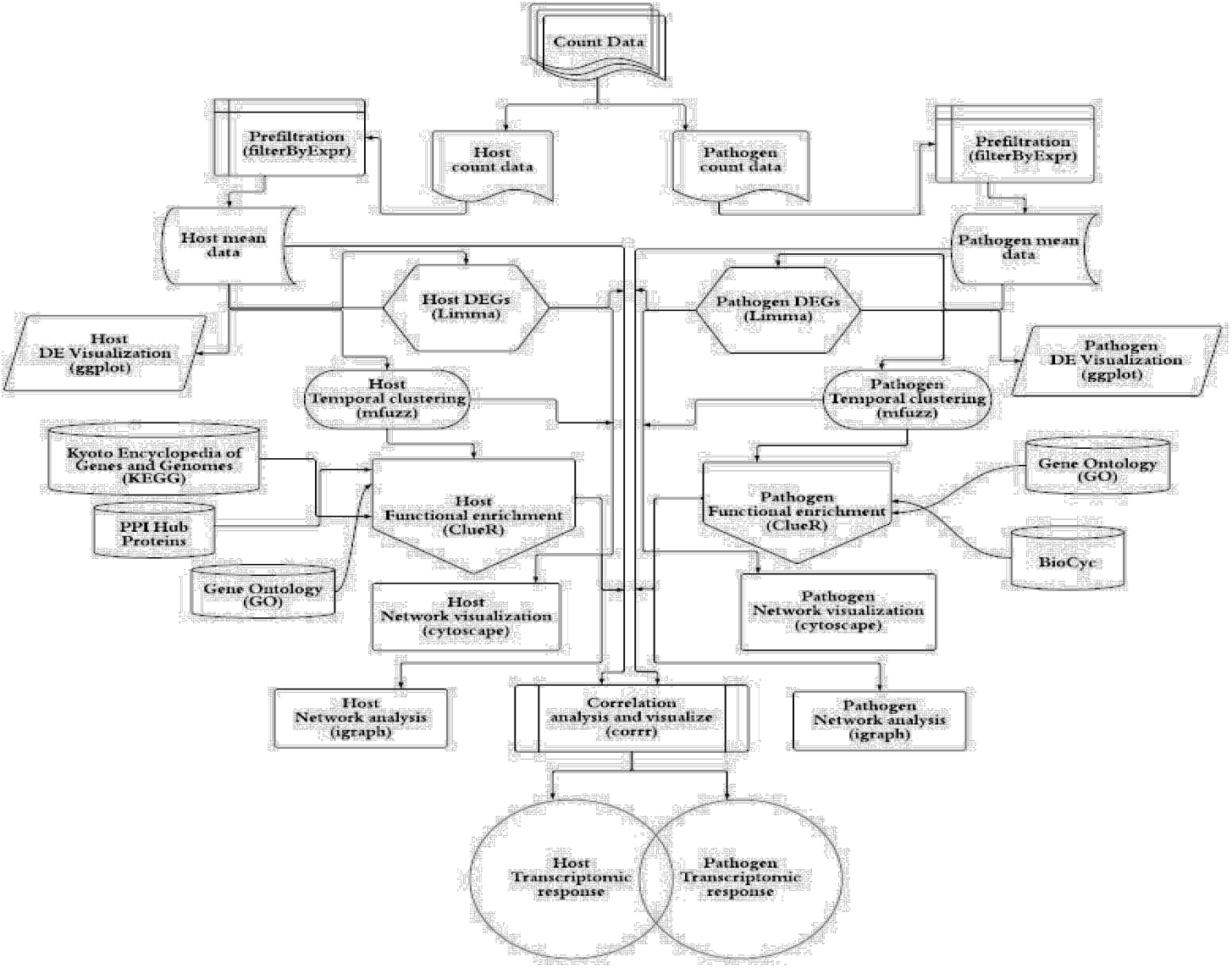
Flowchart of the bioinformatic pipeline for the dRNA-Seq analysis of a host – pathogen interaction. The pipeline starts with data pre-processing and differential gene expression analysis following by Fuzzy clustering to decipher coherent patterns of temporal gene expression profiles. The pathway enrichment analysis was applied using KEGG and GO annotations to identify functions overrepresented by temporal clusters in host and pathogen (based on Fisher’s exact test with hypergeometric null hypothesis). To explore relationship and potential physical or regulatory interactions among differentially expressed genes, the protein-protein interaction (PPI) for both species and regulatory networks for pathogen were retrieved from different datasets. The topological characteristics of the genes were then identified. Additionally, the gene co-expression networks were constructed to infer cross-species gene associations and then used to identify hubs, betweenness centrality, closeness centrality and modularity followed by functional analysis.

The dRNA-seq read count matrices were downloaded from Gene Expression Omnibus (accession number GSE60144) (25) quantifying the expression level of host and pathogen transcripts across six time-points (0 h, 2 h, 4 h, 8 h, 16 h, and 24 h). Gene expression levels in the host and pathogen at different times compared to the start of the experiment (0 h) were evaluated and DE genes (adjusted *p*-value < 0.05 and |log2(fold-change)|>1) were identified. There were 6,248 and 831 DE genes were found in the host and pathogen, respectively, that were differentially expressed at least in one time-point (Supplementary Table). The overlap of upregulated and downregulated DE genes in the host and pathogen is displayed in Figure 2. As can be seen, the number of downregulated genes in the host (Figure 2A) and the pathogen (Figure 2B) were significantly increased at 2h and 24h pi (compared to pre-infection), respectively. A higher number of upregulated genes in both organisms was observed at 24h pi. Venn Diagrams in Figure 2 also present the number of genes dysregulated specific to each time point.

**Figure 2.**
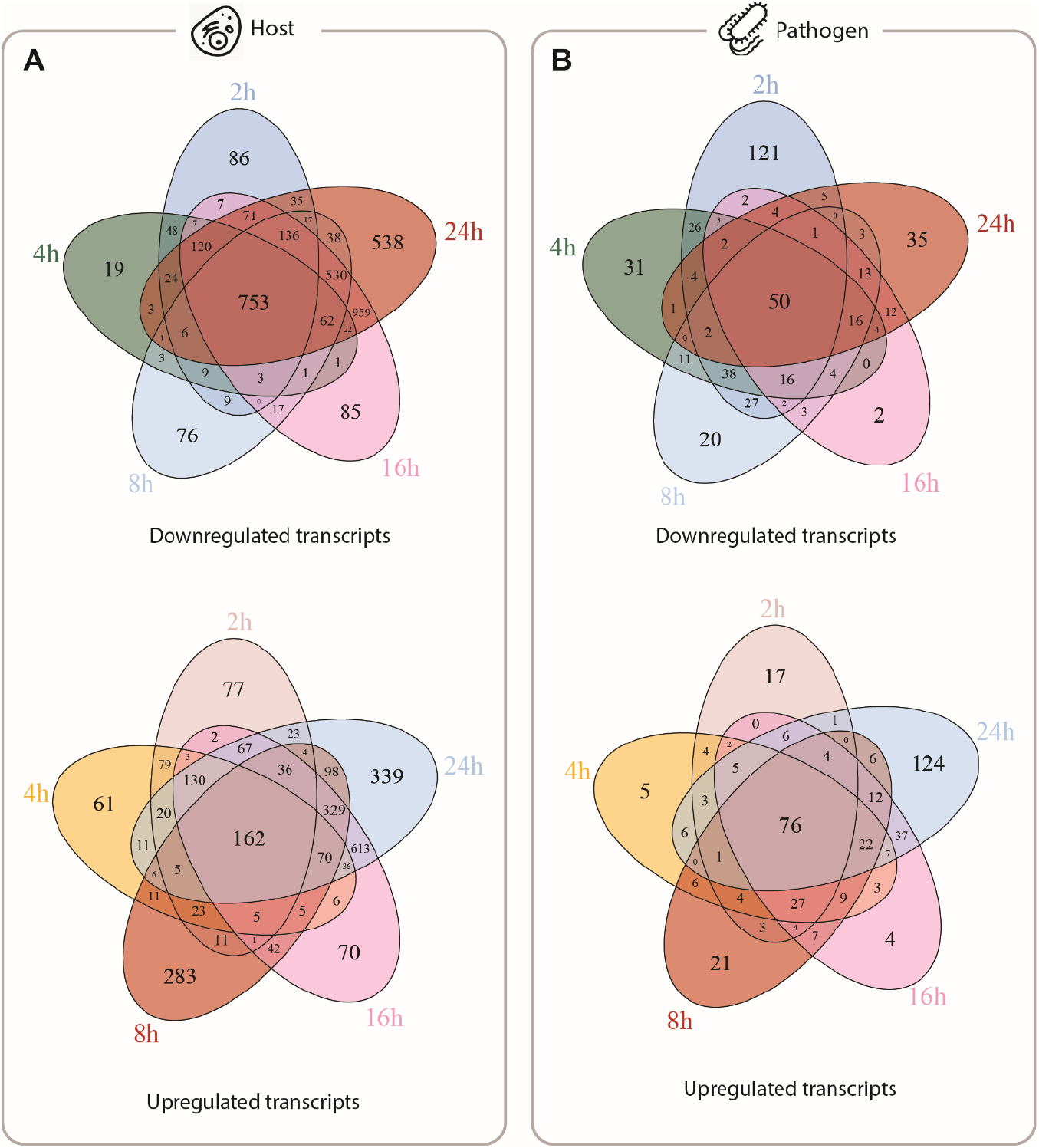
DEGs analysis of host and pathogen. The Venn diagrams indicating numbers of DE genes in the host and pathogen cross five different timepoint compared to 0 h (adjusted *P* < 0.05).

In general, genes with similar expression patterns which form temporally co-expressed clusters have the potential to exhibit similar cellular functions as guided by the “guilt-by-association” assumption (40). Accordingly, fuzzy clustering algorithm which produces overlapping clusters was applied to the 6,248 host and 831 pathogen genes with expression time series of length 6. The optimal number of clusters was 10 and determined as previously explained (see Methods). The membership degree was set to ≥ 0.5 to select genes that tightly follow the average expressional pattern of each cluster and remove genes which do not strongly belong to any cluster.

Accordingly, 4,893 out of 6,248 genes grouped into 10 clusters (Figure 3A) and the most populated expression pattern observed as upregulation and downregulation activities peak at 24 h (pi) in clusters six and ten, respectively (Table 1). On the other hand, the result of pathogen clustering showed 690 out of 831 pathogen genes were grouped into 10 clusters (Figure 3B); the most populated expression pattern observed as an up-regulation and down-regulation activities peak at 24 h (pi) in clusters three and ten, respectively (Table 2).

**Figure 3.**
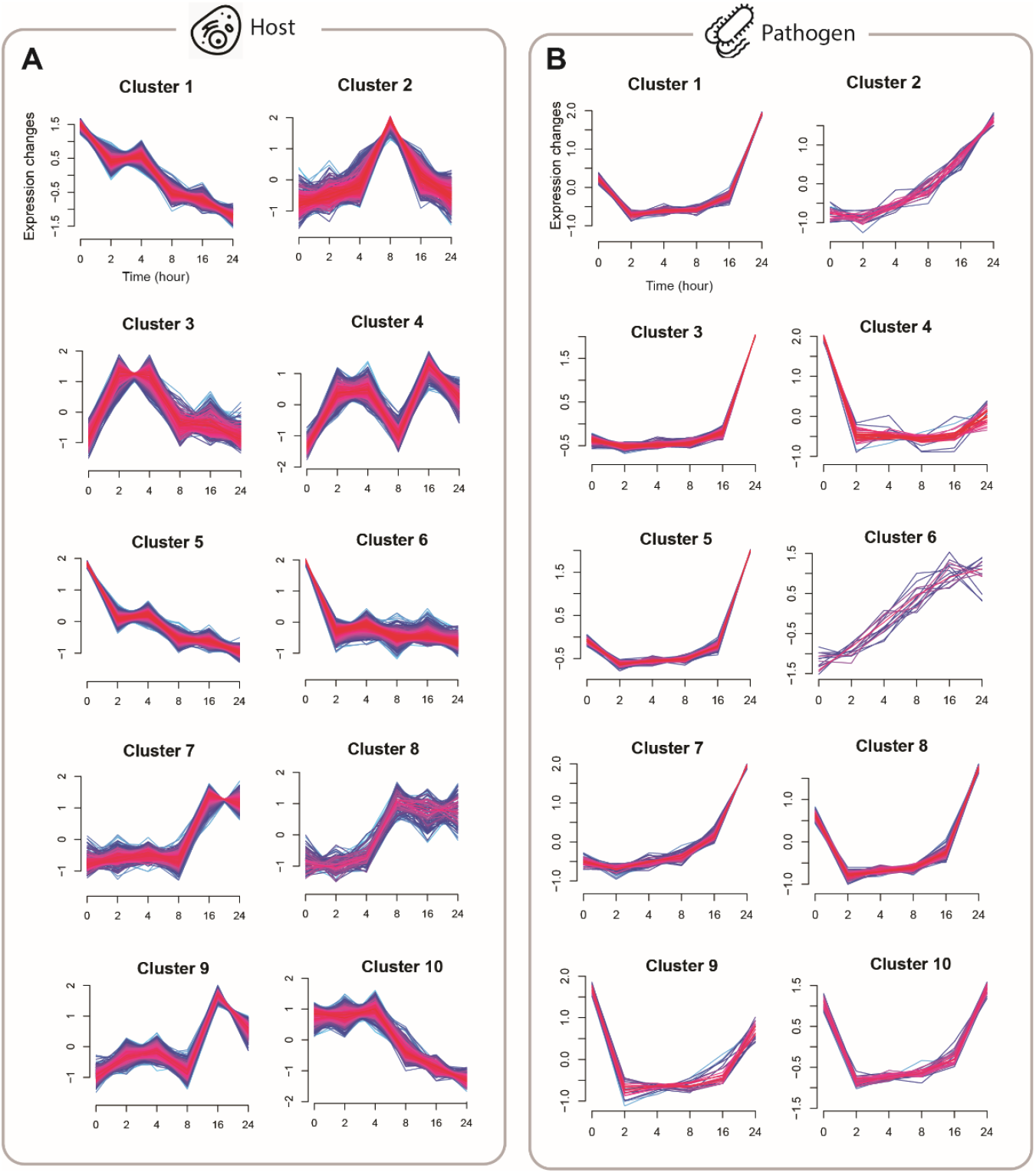
Clusters representing temporal expression patterns of genes differentially-expressed in host and pathogen. The y-axis represents the standardised mean expression at each timepoint.

**Table 1.** Summary of DEGs distribution in the host clusters.

| Timepoint (pi) | Total genes | Upregulated | Downregulated |
| --- | --- | --- | --- |
| 2h | 1646 | 492 | 1154 |
| 4h | 1413 | 449 | 964 |
| 8h | 2164 | 702 | 1462 |
| 16h | 3620 | 1151 | 2469 |
| 24h | 4328 | 1412 | 2916 |

**Table 2.** Summary of DEGs distribution in the pathogen clusters.

| Timepoint (pi) | Total genes | Upregulated | Downregulated |
| --- | --- | --- | --- |
| 2h | 381 | 115 | 122 |
| 4h | 324 | 137 | 187 |
| 8h | 328 | 146 | 182 |
| 16h | 310 | 190 | 120 |
| 24h | 408 | 286 | 266 |

The functional annotation and enrichment analysis on clusters of DE genes using GO terms and KEGG allowed us to obtain functions enriched by temporal clusters of DE genes in host (Supplementary Table) and pathogen (Supplementary Table) during the infection process.

Overall, 20 genes from four pathogen clusters were enriched with two main classes of cellular functions important for pathogenesis (Supplementary Table). The first class was related to the transmembrane transporter activity, electron transport chain, and the biosynthesis of a micronutrient such as biotin that acts as a co-factor for bacterial metabolic activities. The role of biotin biosynthesis and transport functions in bacterial pathogens have been well studied (41–43). Therefore, understanding the transcriptional mechanisms regulating essential elements of biosynthesis and transport will extend our knowledge on bacterial survival and metabolic adaptation during pathogenesis.

The peptidoglycan biosynthetic pathway is a critical process in the bacterial cell (44) and the interplay between peptidoglycan biology and pathogenesis has been previously reported (45). The enrichment analysis has shown that the second class of functions enriched among pathogen DE clusters included the regulation of cell shape and peptidoglycan synthesis that enable bacteria to maintain, modify, and reshape the peptidoglycan layer without risking its essential functions in cell shape, cellular integrity, and intermolecular interactions.

During the initial steps of infection, pathogens manipulate and modify host cell biology to ensure conditions best suited for colonising (46). This process can stimulate the response of the innate immune system (47). GO enrichment analysis has shown that 380 genes across host clusters were enriched with functions which play essential roles in mediating cellular entry of pathogens (*e.g*., epidermal growth factor receptor (EGFR) signalling pathway (48), growth hormone receptor signalling pathway (49), mitochondrial ATP synthesis coupled electron transport (50), protein targeting to endoplasmic reticulum(51)), communication of bacteria with the host environment (*e.g*., positive regulation of establishment of protein localization(52)), control of infection (*e.g*., regulation of protein metabolic process(53)), invading pathogen (*e.g*., response to laminar fluid shear stress (54)), and co-opting host factors (*e.g*., viral gene expression (55)) that can be considered as potential targets in designing antibacterial therapies. Enriched functions were mostly observed at 24h and 4h pi in the host and pathogen, respectively, with mainly downregulated expression (Supplementary Table).

Integration of biological networks, such as protein–protein interactions, has shown to be a useful approach to correlate gene expression changes with changing conditions (56, 57), and can provide information about the most significant connections among classified DE genes in the host and pathogen during early stages of infection. We, therefore, overlaid DE genes across different clusters on protein-protein interaction (PPI) network of the host and pathogen to show how different clusters can possibly cross talk. We also constructed PPI networks among *all* DE genes (not exclusive to those genes appeared in temporal clusters) in the host and pathogen that, in this study, are referred to as the host gene network (HGN) and the pathogen gene network (PGN), respectively.

In order to detect key genes and clusters of genes with functionally enriched pathways during early stages of infection, a gene co-expression network analysis using *igraph* package was performed. Co-expression modules identified by clustering are often large, and therefore, it is important to identify key gene(s) in each module. A widely used approach is to identify highly connected genes in a co-expression network (i.e., hub genes). A hub gene is a gene associated with many other genes in gene networks. Additionally, centrality measures, mainly ‘betweenness centrality’, are often used to identify topologically important genes in a biological network (58). High betweenness centrality of a gene indicates that the node/gene serves as a connector in the network and lies in many shortest paths that connect together different parts of the network (59). Genes with high degree and betweenness centrality may play a key role in biological processes and gene regulation (60) and tend to be highly relevant to the functionality of biological networks. Many biological networks are organised into connected substructures or modules which carry particular functions within the system (61). Hub genes can be identified within a module (rather than the entire network), that is referred to as intra-modular hubs and can be central to a specific function carried by the module. Correlation analysis has shown that hub genes, i.e., nodes with degree > 10 for pathogen and > 100 for host, separately identified in the host and pathogen co-expression networks (129 and 293 hubs in host networks, respectively), are significantly correlated. We also applied network community detection (louvain clustering algorithm) on hub genes and identified four inter-connected modules in negatively correlated (corr < −0.7) (Figure 4A) network and three modules in positively correlated (corr > 0.7) (Figure 4B) network. These modules can potentially be involved in the same or similar biological processes.

**Figure 4.**
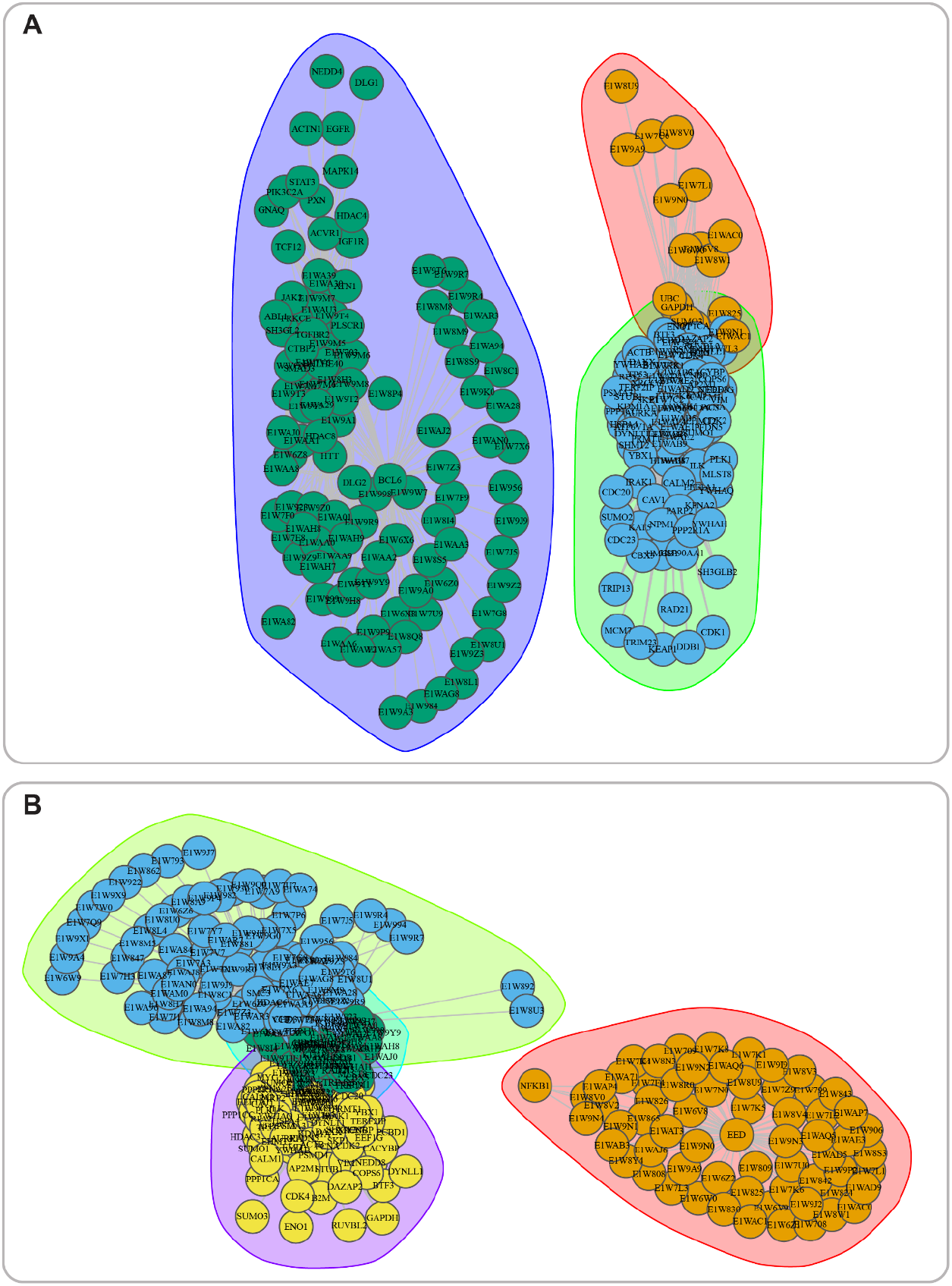
Modules of correlated host and pathogen genes with high degree and betweenness. **A.** Negative correlation, **B.** Positive correlation.

To identify potential cellular functions and diseases associated with genes contained in each module, we performed KEGG pathway, KEGG disease, and GO enrichment analyses on the set of genes contained in each of the negatively and positively correlated modules. Analyses have shown that the host and pathogen gene sets in the negatively and positively corelated modules were enriched with similar as well as host-/pathogen-specific functions (Table 3). For instance, the most frequently enriched functions specific to the host gene set in the first negatively-correlated module were involved in processes underlying the protection against infectious diseases, and the innate immune system which employs germline-encoded pattern-recognition receptors (PRRs) (62, 63) recognition receptors (63, 64). These distinct processes are activated upon detection of the initial natural infection mediated by a pathogen or immunization.

**Table 3.**
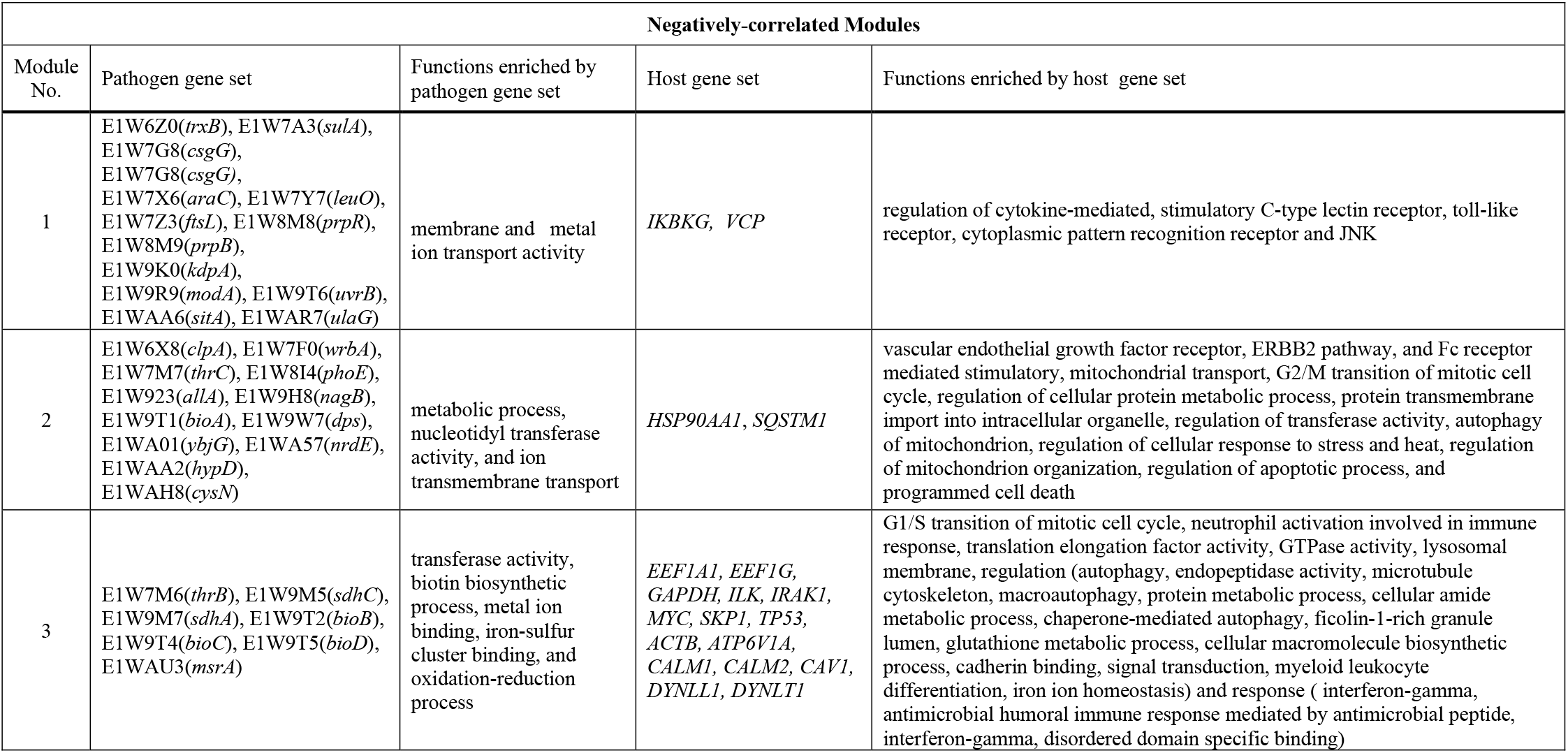
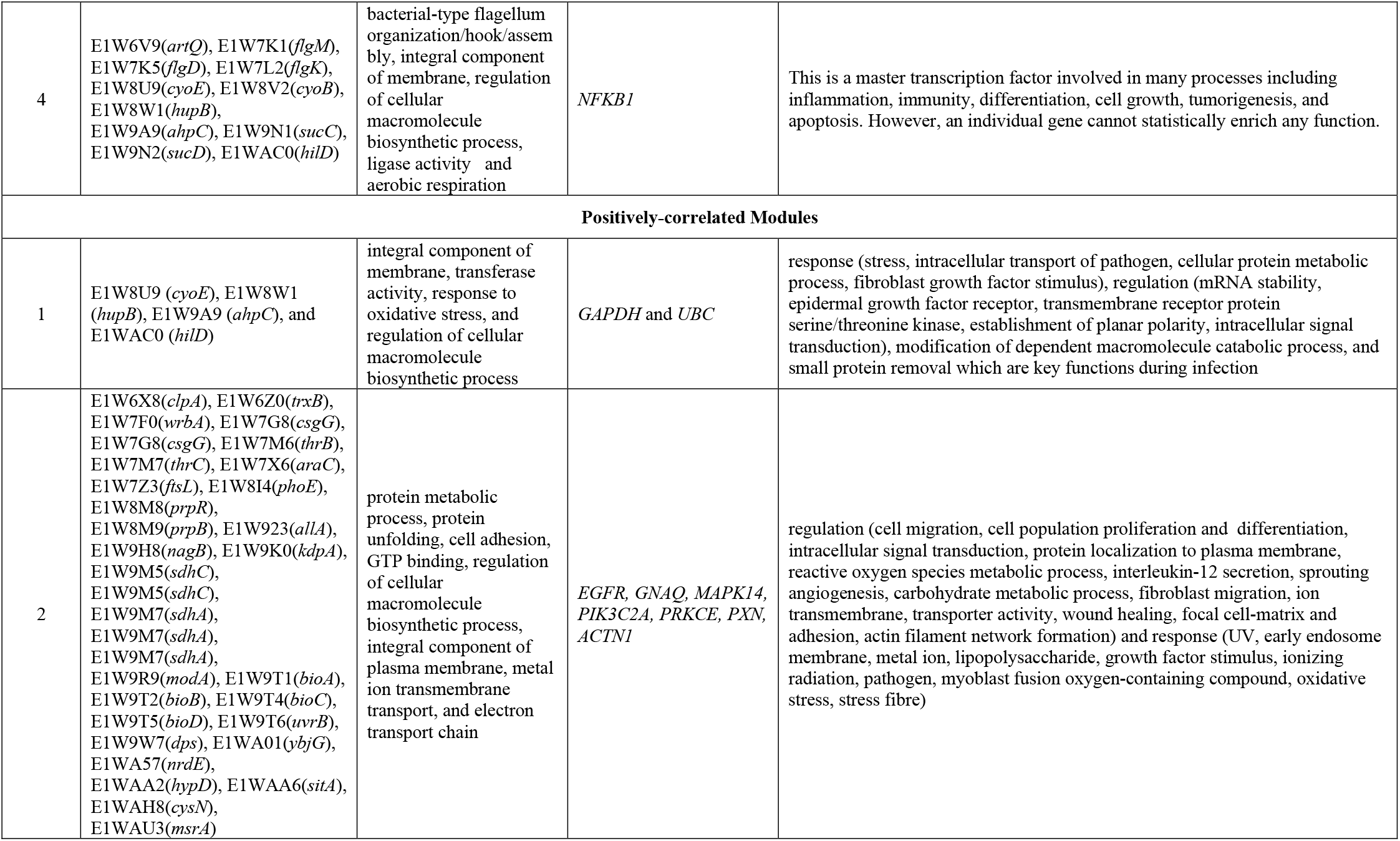
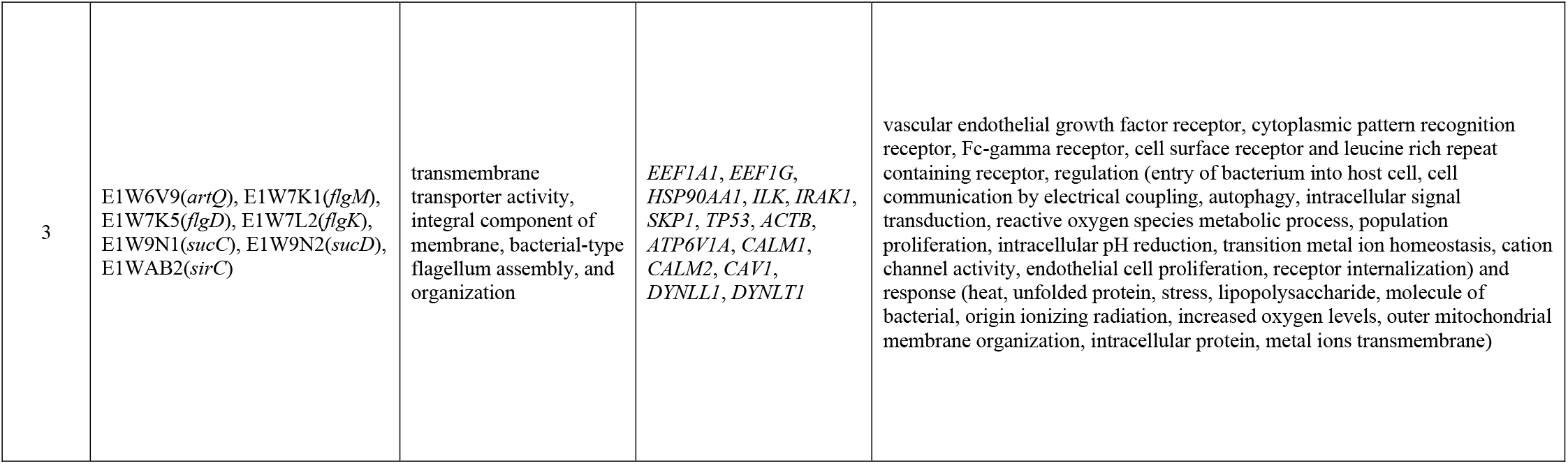
Enrichment analysis (KEGG pathways and Gene ontology) by host and pathogen DE genes in each of positively- and negatively-correlated modules.

The mitochondria metabolic process was the main function that enriched by host genes in the second negatively correlated module. This function has a central role in regulating cellular activities (65) and directly control host stimulated responses against infections (65). Yet, multiple pathogens have the ability to manipulate mitochondrial dynamics and functions as well as cell metabolism and immune responses (66, 67).

The Wnt signalling pathway was a dominant pathway enriched by the host gene set in the third negatively-correlated module. This pathway contributes to the cell cycle control, cytoskeleton reorganization during phagocytosis and cell migration, autophagy, apoptosis, and a number of other inflammation-related events (68). It has also emerged as an integral component of host responses to infection but its function in the context of immune responses is not fully understood (68–70). Recent studies have shown that this pathway can control host processes related to bacterial infection; however, pathogens have evolved strategies to manipulate Wnt-associated processes in order to enhance infection and survival within the human host (68–70). The mitogen-activated protein kinase (MAPK) signalling pathway was the second most enriched pathway by this set of genes. This pathway plays a central role in host-pathogen complex interactions and is pivotal for triggering host immune response against pathogens (71, 72).

In the fourth negatively-correlated module, lipid droplets (LDs) were enriched by the host gene set, which are the chock point in the conflict between host and pathogen for nutrients (73, 74). Previous studies on the nutritional status of the infected host have discussed the malfunctioning or lack of LDs as a key player in the regulation of systemic innate immunity (74). On the other hand, bacteria can uncouple LDs from mitochondria and increase host-pathogen contacts by reducing fatty acid metabolism (75). Yet, the nature of distinct junction between the front line of the body defence (LDs) and bacteria is not well unknown (74). Furthermore, recent studies have revealed pathogen ability in the manipulation of the host membrane to facilitate pathogenic entry and recruitment of specific host lipids for the maintenance of favourable intracellular niche to augment their survival and proliferation (73, 74, 76).

In the first positively-correlated module, the host genes enriched the Notch signalling pathway that is critically involved in developmental processes and regulating immune responses during infections (77). Two main signalling pathways enriched by the host gene set in the second positive module included G protein-coupled receptors and epidermal growth factor receptors (EGFR). These are involved in the pathogen pathophysiological processes (78, 79) and in mediating cellular entry of numerous pathogen cells (48), respectively. Generally, the functions related to this module have a role in immune pathogenesis by activities of innate and adaptive immune systems.

Leucine-rich repeats (LRRs) were enriched by the host gene set in the third positively-correlated module. The leucine-rich repeat domain presents in several innate immune receptors, where it is frequently responsible for sensing danger signals and specific pathogen-associated molecules. It will be active the innate immune system as a host defence systems, where they sense specific pathogen-associated molecules (80). Fc-gamma and cell surface receptors were the second signalling pathways which were enriched by the host gene set. These receptors are an essential component in many immune system functions such as the phagocytosis and have the ability to stimulate innate immune responses, releasing inflammatory mediators (46, 81).

Vascular endothelial growth factor (VEGF), also enriched in this module, is known to play crucial roles in endothelial cell proliferation, migration, angiogenesis, vascular permeability, inhibition of apoptosis, and pathogen infection. This pathway is also involved in the interaction between pathogens and host cells as pathogens are able to encode VEGF homologs which bind to the VGFR receptor (VEGFR-2) of the host cells and then contribute to the infection process (82).

Another interesting pathway was Pattern Recognition Receptors (PRRs) which are protein found in host and play role in adaptive immunity as part of the innate immune system. (83) The function related to this module have a role in the protein modification process, cell junction assembly and substrate adhesion-dependent cell spreading. Most functions in this module are involved in infection development and stress responses.

Interestingly, an important function frequently enriched by both host and pathogen gene sets across different modules was related to the regulation of metal ions transmembrane transporters. Metal ions are essential for living organisms and play a vital role in the regulation of both microbial virulence and host immune responses. Recent studies have shown a role for metal ions in protection of the host against infections (84–87). Therefore, it could be beneficial for the invading pathogen to express a variety of such genes in order to steal metal ions from the host.

In both groups (positively- and negatively-correlated modules), KEGG enrichment analyses of the host gene set showed similarities to previous observations during various Gram-negative infections (*Legionella, Salmonella, Shigella, Yersinia, Pertussis, Helicobacter pylori*).

Overall, we identified co-expression modules which are potentially involved in specific biological processes during early stages of the *Salmonella* infection. We then further investigated the inter-module relationships to identify genes that are key in interconnecting different modules. Accordingly, we first selected host-pathogen corelated genes to further stringent the subsequent analyses to modules whose constituent host/pathogen genes have a very similar expressional pattern during infection. This resulted 54 genes with positive or negative correlations (Figure 5) with corr < −0.9 or corr > 0.9 forming six negative (Figure 6A) and positive modules (Figure 6B). We computed network-based metrics on genes within these modules and selected genes connecting different modules with high betweenness (≥40) and degree (≥10) as key inert-module connectors. The analyses uncovered the role of E1W9K0 (*kdpA*), *CAV1*, E1WA57 (*nrdE*) and *SKP1* in interactions between four negative correlated modules (Supplementary Table). Additionally, E1WAC0 (*hilD*), *ATP6V1A, ILK*, E1W6Z0 (*trxB*), *E1W8W1*(*hupB*), and E1W9A9 (*ahpC*) were identified as key genes to interconnect positive correlated modules (Supplementary Table). Further, DE analyses have shown downregulation activities of *EEF1A1*, E1W8W1 (*hupB*), and *CALM2* switching off the expression of E1W6Z0 (*trxB*) in the module one. Furthermore, the downregulation of *ILK* switches off the expression of E1W9H8 (*nagB*) in the module three. Enrichment analyses have shown pathogen genes involved with oxidation-reduction process and potassium ion transport. Host genes mainly related to regulation of cellular response to inflammation, cellular iron ion homeostasis, and regulation of epithelial cell differentiation. Together, these observations support interactions of these genes in the early stage of infection.

**Figure 5.**
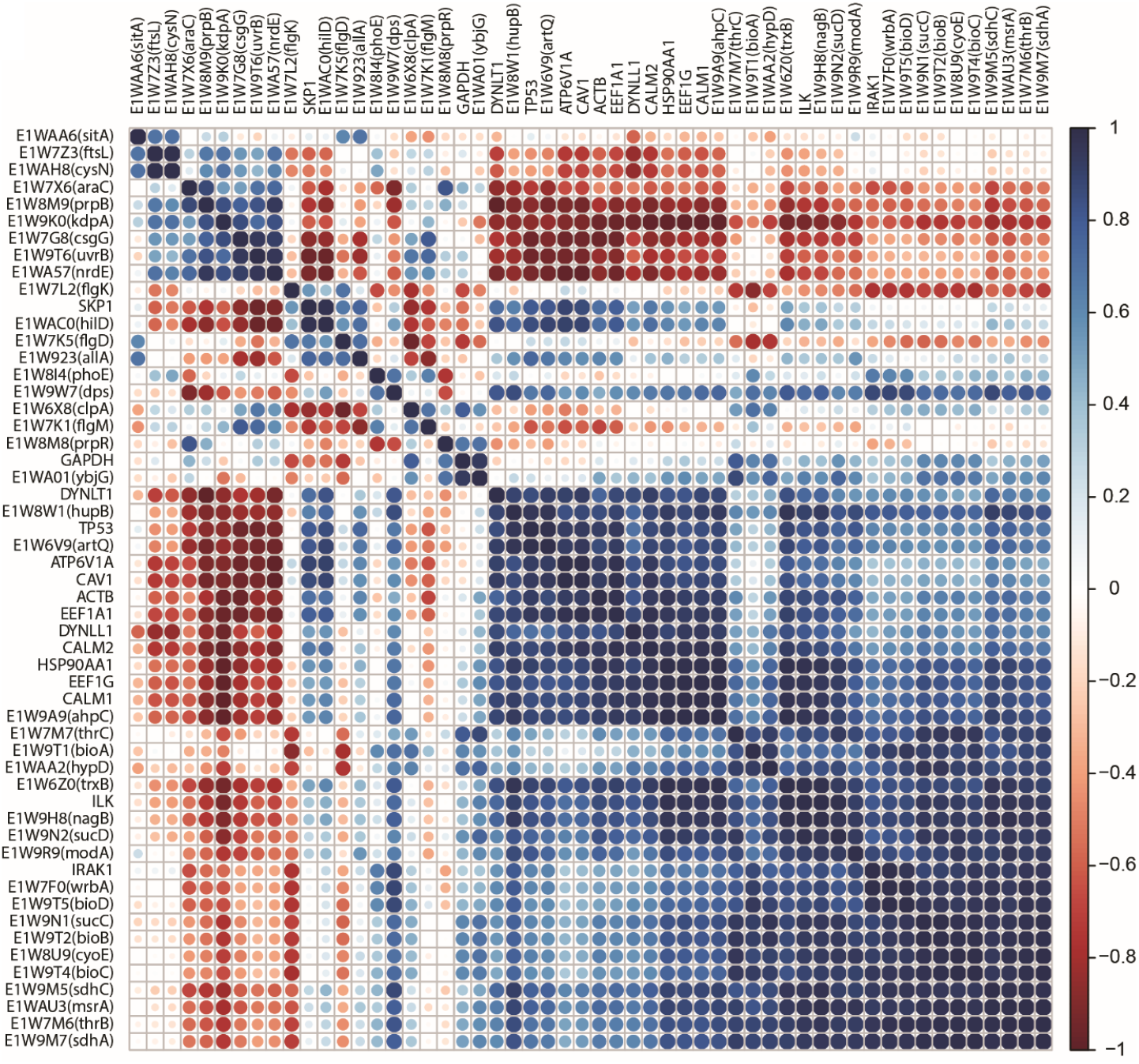
Correlation matrix representing associations within and between host and pathogen DE genes (corr < −0.9 or > 0.9).

**Figure 6.**
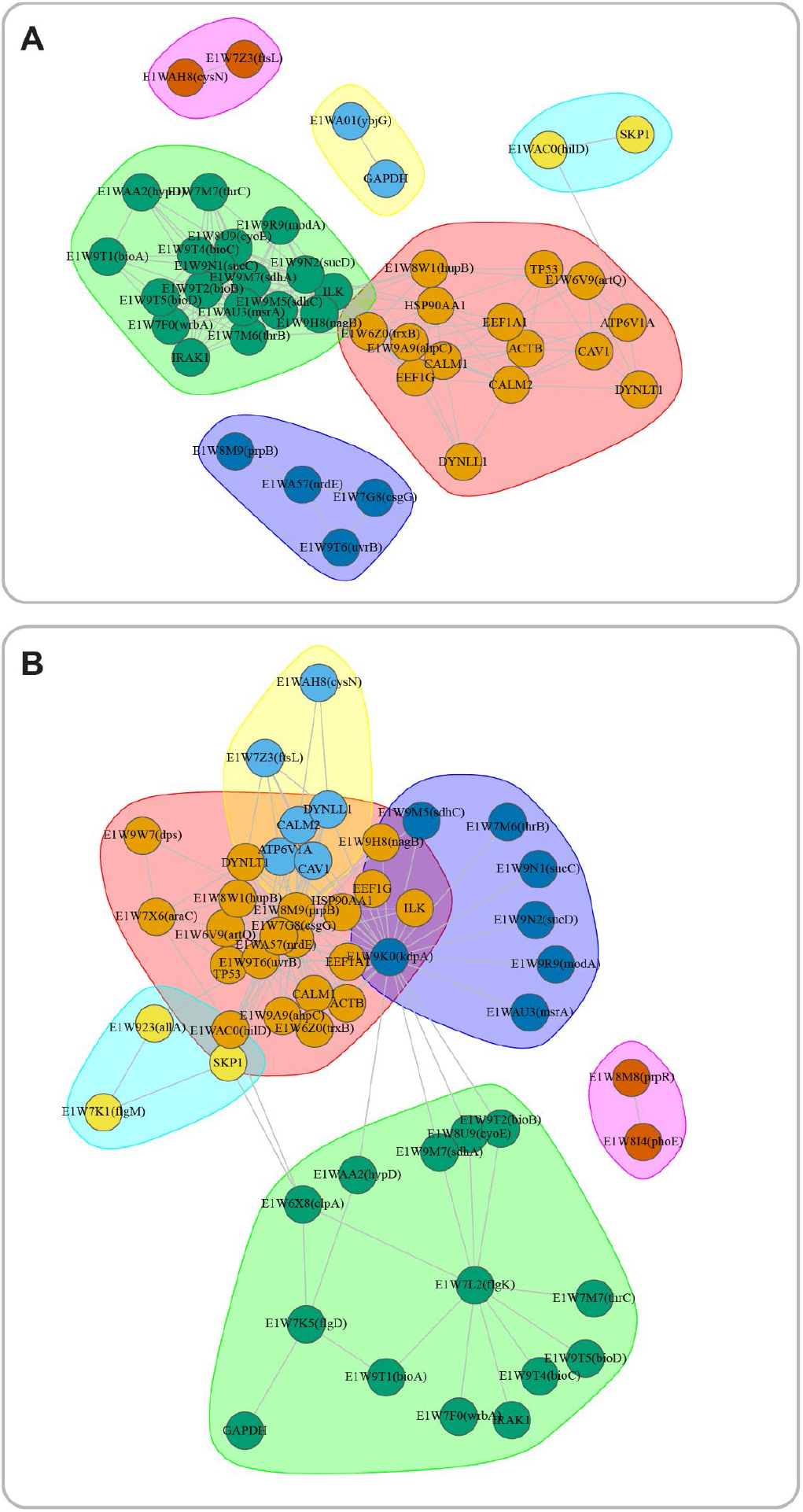
Highly correlated Modules in the host and pathogen genes set. **A.** Negative correlation, **B.** Positive correlation.

## Conclusion

Here, we present the dRNASb pipeline for the network-level analyses of dRNA-Seq data. The pipeline was show cased on a *Salmonella enterica serovar* Typhimurium-infected HeLa cell model, with a detailed description of analyses performed, and results achieved. The proposed network-based analyses provide complementary information on infection processes not obtainable otherwise. In a nutshell, combining statistical analyses with network-level and functional analyses, as presented in this study, can elucidate the interplay between host and pathogen during bacterial infection and provide valuable information on which genes in the host or pathogen can be potential molecular targets for therapeutic interventions. Development of such pipelines can enhance system-level understanding of host-pathogen transcriptomic dynamics and can aid in the identification of key genes or functional interpretation of poorly annotated genes. In the future, dRNASb can be further enhanced by inclusion of machine learning models to further integrate complementary information for accurate prediction of host-pathogen interactions during the infection process.

## Data and Code Availability

The data set used to showcase the pipeline is available in NCBI’s Gene Expression Omnibus under accession number GSE60144. The pipeline code is available at GitHub repository https://github.com/VafaeeLab/dRNASb.

## Author Statements

### Author Contribution

Conceptualisation: FV, MD; Methodology: MD, FV; Software: AV, MD; Validation: MD, NCR, AP; Formal Analysis: MD, FV, FK, DM, AFT, KV, AA; Investigation: FV, MD; Resources: FV; Data Curation: MD; Writing – Original Draft Preparation: MD, FV; Writing – Review and Editing: All Authors; Visualisation: MD; Supervision: FV; Funding: FV, NCR.

### Conflicts of interest

Authors declare no conflicts of interest.

### Funding information

NCR is supported by a Cancer Institute NSW Early Career Fellowship (2019/ECF1082) and a UNSW Scientia Fellowship. Otherwise, not externally funded.

## Acknowledgements

We thank Dr Susan Corley, Dr Sara Loo, and Dr Xabier Vázquez-Campos for useful discussions and support.

## Notes

### Competing Interest Statement

The authors have declared no competing interest.

